# Identification of viral-mediated pathogenic mechanisms in neurodegenerative diseases using network-based approaches

**DOI:** 10.1101/2020.12.21.423742

**Authors:** Anna Onisiforou, George M. Spyrou

## Abstract

During the course of a viral infection, virus-host protein-protein interactions (PPIs) play a critical role in allowing viruses to evade host immune responses, replicate and hence survive within the host. These interspecies molecular interactions can lead to viral-mediated perturbations of the human interactome causing the generation of various complex diseases, from cancer to neurodegenerative diseases (NDs). There are evidences suggesting that viral-mediated perturbations are a possible pathogenic aetiology in several NDs such as Amyloid Later Sclerosis, Parkinson’s disease, Alzheimer’s disease and Multiple Sclerosis (MS), as they can cause degeneration of neurons via both direct and/or indirect actions. These diseases share several common pathological mechanisms, as well as unique disease mechanisms that reflect disease phenotype. NDs are chronic degenerative diseases of the central nervous system and current therapeutic approaches provide only mild symptomatic relief rather than treating the disease at heart, therefore there is unmet need for the discovery of novel therapeutic targets and pharmacotherapies. In this paper we initially review databases and tools that can be utilized to investigate viral-mediated perturbations in complex NDs using network-based analysis by examining the interaction between the ND-related PPI disease networks and the virus-host PPI network. Afterwards we present our integrative network-based bioinformatics approach that accounts for pathogen-genes-disease related PPIs with the aim to identify viral-mediated pathogenic mechanisms focusing in MS disease. We identified 7 high centrality nodes that can act as disease communicator nodes and exert systemic effects in the MS enriched KEGG pathways network. In addition, we identified 12 KEGG pathways targeted by 67 viral proteins from 8 viral species that might exert viral-mediated pathogenic mechanisms in MS by interacting with the disease communicator nodes. Finally, our analysis highlighted the Th17 differentiation pathway, a hub-bottleneck disease communicator node and part of the 12 underlined KEGG pathways, as a key viral-mediated pathogenic mechanism and a possible therapeutic target for MS disease.

## Introduction

Neurodegenerative diseases (NDs) are chronic degenerative diseases of the central nervous system (CNS) and currently there are no effective pharmacotherapies for their treatment, thus highlighting the need for novel therapeutic interventions. Although several aetiologies have been identified to contribute to their development, the exact underlying mechanisms are still unclear. Viral infections have been suggested as a possible pathogenic aetiology of NDs as viruses can cause degeneration of neurons via both direct and/or indirect actions and several viruses have being associated with NDs as described in Table 1 [1–3]. Viruses can induce neuronal dysfunction directly through their cytolytic effects, and indirectly via various mechanisms such as by expressing viral genes that interfere with the host’s immune system and cellular processes, via bystander inflammatory reactions, or by inducing apoptosis [4,5].

**Table 1:** Viruses associated with NDs

| <u>Neurodegenerative Disorder</u> | <u>Associated Viruses</u> | <u>References</u> |
| --- | --- | --- |
| **Alzheimer's Disease** | Herpes simplex virus 1 (HSV-1), Human Immunodeficiency Virus (HIV), Herpes Simplex Virus 2 (HSV-2), Human Cytomegalovirus (HCMV/HHV-3), Epstein-Barr Virus (EBV/HHV-4), Varicella-Zoster Virus (VZV/HHV-5), Human Herpesvirus 6 (HHV-6), Human Herpesvirus 7 (HHV-7), Hepatitis C Virus (HCV) | [1,2,12–15] |
| **Multiple Sclerosis** | EBV, HHV-6, HSV-1, Measles *Morbillivirus* Virus, Rubella Virus, VZV, Human Endogenous Retrovirus W (HERV-W), Human T-cell Leukemia Virus type 1 (HTLV-1), John Cunningham Virus (JCV) | [16–25] |
| **Parkinson's Disease** | West Nile Virus, Japanese Encephalitis B Virus, Saint Louis Encephalitis Virus, HIV, HSV, EBV, HCMV, VZV, Influenza Virus Type A, Measles *Morbillivirus* virus, Coxsackie virus, Polio Virus | [1,26–29] |
| **Amyotrophic Lateral Sclerosis** | HIV, HTLV-1, Human Endogenous Retrovirus K (HERV-K), Enteroviruses (Poliovirus, Coxsackievirus, Echovirus, Enterovirus-A71 and Enterovirus-D68) | [1,3,30,31] |

Viruses are intracellular obligate parasites that lack their own replication machinery, thus to ensure their survival and reproduction they rely on the host, and via physical interactions they manipulate and exploit the host’s cellular machinery [6,7]. In addition, through the evolutionary process, viruses have developed an array of adaptive immune evasion strategies including interfering with antigen presentation and mimicking immune processes [5]. Several evidence indicates that neuroinflammation is involved in the progression of NDs, by directly or indirectly contributing to neuronal loss, however whether the immune system participates in the initiation of these diseases remains undetermined [1,8]. Immune system activation within the CNS can also be elicited by viral infections, either by neurotropic viruses that infect the CNS or by viruses that infect peripheral tissues and can cause a strong inflammatory response, resulting in the infiltration of peripheral leukocytes into the CNS and the subsequent activation of resident microglia cells [9]. Infection by certain neurotropic viruses can also lead to the formation of neuropathological and/or immunopathological lesions with the pattern of the resulting lesions having similar distribution as that of the common NDs [10,11]. It is therefore possible that viruses may be directly or indirectly involved in the development of NDs, by either causing dysregulation of the immune system and/or by interfering with the host’s cellular and immune system components.

However, despite the association of viral infections with NDs, it is highly unlikely that viruses are solely responsible for the development and/or progression of NDs, as the prevalence of viruses that are linked to NDs is very common in the general population, whereas the rate of individuals with NDs is significantly lower [4]. Therefore, it is unlikely that viral infections are the self-determining factor in the pathogenesis of NDs, and other factors particularly genetic susceptibility or environmental factors could contribute in association with viral infections. Genetic susceptibility is not only associated with NDs, but is also an important factor that determines how a person’s immune system responds to a pathogen and how efficiently it can eliminate acute viral infections or control chronic viral infections [4,8]. Therefore, the individual’s genome determines the genetic sensitivity of the immune system towards different pathogens. However, genetic sensitivity to infections does not imply that there is a general weak immune surveillance towards pathogens [4], but rather that the individual’s “immunogenome” is more susceptible towards specific pathogens. The “immunogenome” can change virus-host symbiosis and predispose an individual to certain viral-mediated immune associated diseases by various mechanism such as altering immune response to specific viruses, shift viral cell tropism and change neurovirulence [32–34]. Therefore, understanding the interaction between viruses, genes and environment is not only essential for elucidating the role of viruses in the pathogenesis of NDs, but can also aid to determine how genetic predisposition can lead to NDs (Figure 1).

**Figure 1:**
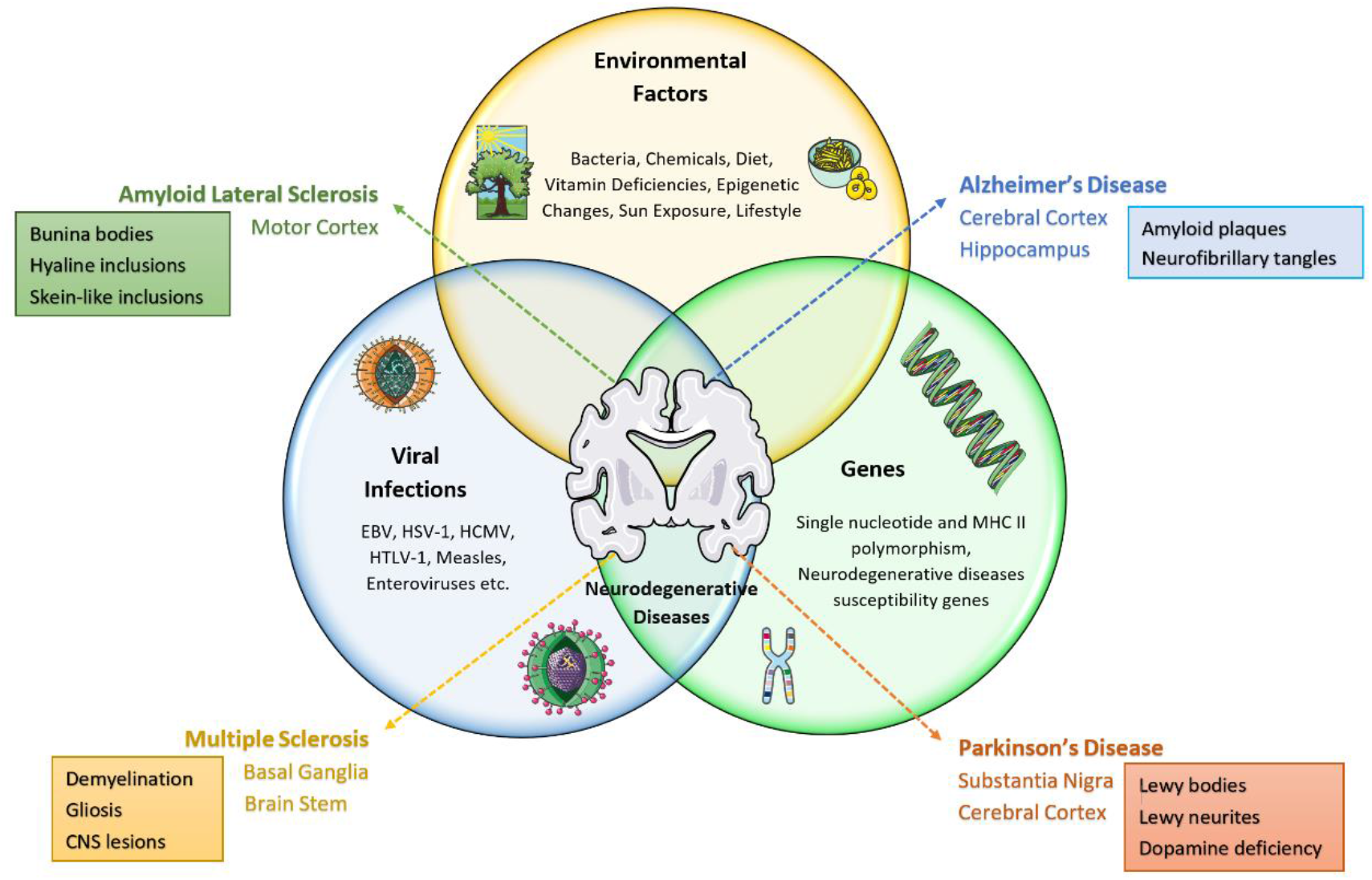
A schematic representation of the multifactorial origin of a group of pathologically distinct but related neurological diseases. NDs are complex disorders that are caused by the interaction of environmental factors, viral infections and genetic susceptibility. The combination of this factors contributes to the clinical heterogenicity and histopathological diversity of the central nervous lesion that characterizes these diseases, as they differ in the subset of neurons and anatomical structures that are affected, and have both common but also distinct pathological abnormalities. Figure contains illustrations obtained from Servier medical art (https://smart.servier.com/), provided free and licensed under the Creative Commons Attribution 3.0 Unported License.

## INTEGRATION OF VIRUS-HOST PPIs AND HOST PPIs IN NDs INTERACTOMES FOR THE INVESTIGATION OF VIRUS-MEDIATED PATHOGENESIS

During the course of a viral infection, virus-host PPIs are a key infection mechanism that allows viruses to evade host immune responses, replicate and hence survive within the host. These molecular interactions can lead to the dysregulation of normal biological processes within the host, causing numerous diseases from cancer to NDs [35]. These interspecies PPIs can be represented as a network, where the nodes represent the proteins and the edges their interactions [36]. This physical interaction network between viral proteins and host’s cellular targets can be used to provide further insights in the disease etiology [37]. Understanding pathogen-host PPIs might enable the identification of mechanisms that directly or indirectly lead to the development of certain diseases.

Human PPI networks exhibit a scale free behavior, where the majority of proteins in the network have few connections to other proteins whereas few ones, termed “hubs”, are connected with multiple proteins [38]. Viral families can be classified based on their genome, DNA or RNA, and comparative interactome analysis of DNA versus RNA viruses human PPIs revealed that the DNA viruses-human PPIs network follows a scale free behavior, whereas the RNA viruses-human network does not follow a scale free behavior [39]. In addition, unlike the cooperative nature of the evolutionary principles that govern PPIs of the host’s biological processes, the evolutionary trajectories of PPIs in the virus-host interaction network are antagonistic, as there is a constant competition between virus and host interactions where a change in a protein of the virus might result in a reciprocal counterchange in a host’s protein and vice versa [40,41]. This constant arm race of evolutionary actions and counteractions between virus-host processes determines the evolutionary nature of the virus-host PPIs network, which is not static and hence its topology changes over time.

Topological analysis studies of virus-human PPIs networks revealed several important characteristics about the interaction of viral proteins with human targets. Viral proteins have a preference to target human proteins that are hubs, have high betweenness (bottlenecks), are articulation points and belong to rich clubs [42–45]. Hub proteins play an essential role in maintaining the connectivity of the network. The removal of hub nodes causes network failure, a property known as lethality-centrality, whereas the deletion of a random node with few connections does not influence the topology of the network [46,47]. Betweenness centrality enables the identification of nodes that act as bridges (bottlenecks) between other nodes in the network and hence are important in influencing the communication between nodes within the network [48]. Viruses through the evolutionary processes have evolved strategies that allow them to efficiently adapt to the scale free nature of the human interactome and via targeted attacks interact with essential proteins (hubs, bottlenecks) that are highly connected, allowing them to influence multiple functions and pathways simultaneously [49].

Viral-mediated perturbations of the host’s interactome that lead to the generation of complex diseases involves the interaction of viral proteins with the host’s PPIs network, therefore merging the virus-human PPIs and the human disease PPIs networks can enable to understand viral-induced pathogenesis [35,37,49,50]. In the context of NDs, the virus-host PPIs network can be merged with the ND-related PPIs networks, creating an integrated virus-host-NDs PPIs network allowing to identify viral-mediated pathogenic mechanisms in NDs (see Figure 2 for a schematic representation).

**Figure 2:**
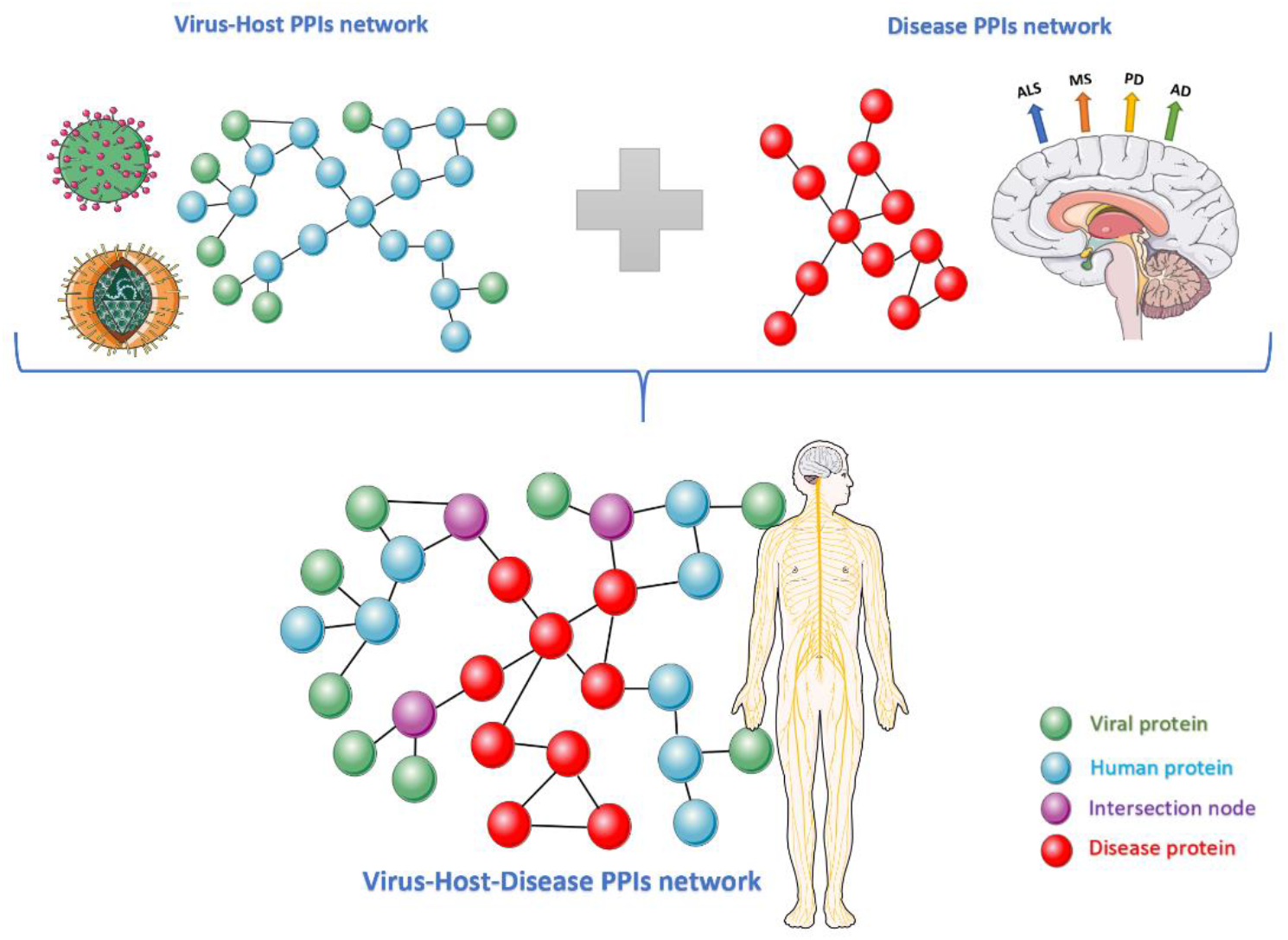
Schematic representation of the construction of the integrated virus-host-NDs PPIs network, where the virus-host PPIs network is merged with a ND-related PPIs network. Figure contains illustrations obtained from Servier medical art (https://smart.servier.com/), provided free and licensed under the Creative Commons Attribution 3.0 Unported License.

## COMPUTATIONAL ANALYSIS OF VIRUS-HOST-DISEASE PPIs USING A KNOWLEDGE BASED APPROACH

The analysis of virus-host-disease interactions can involve several approaches with the most commonly used bioinformatics pipeline approach involving the following steps: (1) reconstruction and visualization of the virus-host-disease PPI network, (2) topological network analysis and (3) Gene ontology (GO) [51] and pathway enrichment analysis. There are several reviews that have extensively described databases and tools that can be implemented to investigate this central steps [52,53]. Figure 3, illustrates these main steps and summarizes some of the recourses that can be utilized at each step for the investigation of virus-host-disease PPIs. Computational analysis using the networks topological properties allows to identify important interactions within the network, with the most commonly used for the investigation of virus-host interactions being: node degree, degree centrality, betweenness centrality, shortest path and modularity score [43,44,48,54,55]. On the other hand, GO and pathway enrichment analysis has been used in several studies to identify common and unique infection strategies, as well as pathogen-mediated pathogenic mechanisms that may lead to disease [39,44,55–57].

**Figure 3:**
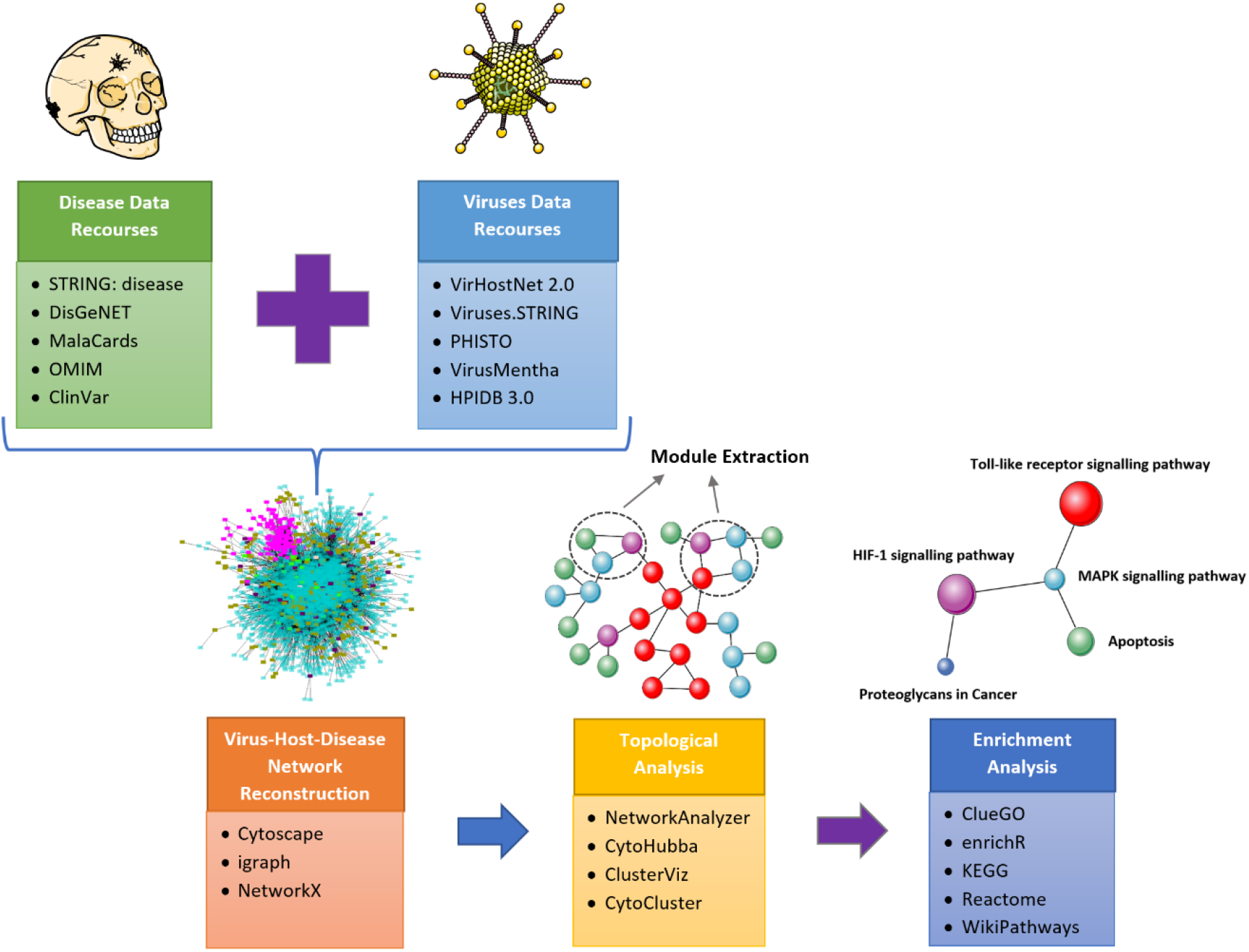
Illustration of the most commonly used approach for the investigation of virus - host - disease PPIs highlighting recourses and tools that can be utilized at each step. The first step involves the collection of data, virus-host PPIs can be collected from several databases such as VirHostNet 2.0 [110], Viruses.STRING [111], PHISTO [80], VirusMentha [112] and HPIDB 3.0 [113]. Disease data can be collected from databases such as the *STRING: disease* query app in Cytoscape [77], MalaCards [74], DisGeNET [75], OMIM [114] and ClinVar [115]. The second step involves the visualization of the integrated network which can be performed with Cytoscape [81], the igraph package in python and R [116] and the NetworkX package in python [117]. The third step involves network topological analysis that can be performed with several plugins offered by Cytoscape such as NetworkAnalyzer [118] and CytoHubba [89], as well as clustering analysis apps, such as CytoCluster [119] and ClusterViz [120]. The final step involves enrichment analysis that can be performed either on the whole network or on subnetworks using the ClueGO [83] plugin in Cytoscape [121] and enrichR [122] an R interface to the Enrichr database [123,124]. Pathway enrichment analysis can be performed form several databases including the Kyoto Encyclopedia of Genes and Genomes (KEGG) [84], Reactome [125] and WikiPathways [126]. Figure contains illustrations obtained from Servier medical art (https://smart.servier.com/), provided free and licensed under the Creative Commons Attribution 3.0 Unported License.

Although several studies have investigated the role of viral-mediated perturbations in the generation of human diseases based on PPIs networks [35,37,42,54] there is lack of studies that focus specifically on viral-induced pathogenic mechanisms in the generation of NDs. A recent study by [58] has investigated the mechanisms of viral-induced pathogenesis in NDs, but at the transcriptomic level by comparing the gene expression profiles of AD or PD patients with three viral infection datasets and was able to identify shared pathways between these NDs and viruses. However, despite the lack of studies that explore virus-NDs PPIs there are several studies that have investigated NDs based on PPIs networks. Although NDs encompass a large group of diseases that share as a common pathological component the degeneration of neurons, there is heterogenicity in the subset of neurons, anatomical structures and pathological abnormalities that are affected in each of these diseases [59,60]. Nonetheless, these diseases share several common pathological mechanisms including protein misfolding and aggregation [61], apoptosis [62], impaired bioenergetics [63], neuroinflammation [64], oxidative stress and decreased antioxidant activity [65].

Therefore, NDs are a group of pathologically distinct but related diseases and two different directions have been used to investigate NDs using PPI networks by either focusing on a specific ND or analyzing a group of NDs, with the majority considering a specific disease [66–68]. The few studies that have investigated a group of NDs have primarily concentrated in identifying common molecular mechanisms by identifying direct protein/genes commonalities and pathways among NDs and indirect network relationships between NDs via topological association by identifying common modules in these diseases [69–72]. NeuroDNet is one of the few available specific NDs databases that contains information for genes and SNPs associated with 13 NDs and also allows the reconstruction and analysis of these NDs networks using PPI, regulatory and Boolean networks, however it was last updated in 2016 [73]. The main approaches utilized for the reconstruction of NDs PPI networks are by using disease genes obtained from gene to disease association databases, such as MalaCards [74] and DisGeNET [75], or using gene expression data from NDs patients obtained from the NCBI GEO database [76] and then converting these genes into the corresponding protein products. Another approach is to use the *STRING: disease* query app in Cytoscape [77] which provides directly disease associated proteins, that are obtained from the DISEASES database [78] that collects gene to disease associations from several types of sources and the different types of evidence are unified by assigning them a confidence score. In the next part of our paper we aim to describe a simple methodology that integrates approaches for the investigation of NDs based on PPIs and elements from the investigation of virus-host PPI networks. Our approach aims to incorporate knowledge-based information about a group of NDs with the aim to focus in the molecular pathology of viral-mediated perturbations in MS disease.

## THE CASE OF MULTIPLE SCLEROSIS

Here we present our integrative network-based bioinformatics pipeline approach, illustrated in Figure 4, with the aim to identify possible mechanisms of viral-induced pathogenesis in MS disease. Our pipeline includes the following approaches: (1) reconstruction and visualization of the integrated virus-host-MS PPIs network, (2) topological and knowledge-based subnetwork identification, (3) KEGG pathway enrichment analysis, (4) a filtering process of the enriched results, (5) comparison and identification of direct common pathological pathways in NDs, (6) construction of the MS enriched KEGG pathway-to-pathway network, (7) topological analysis and (8) similarity calculations and clustering analysis.

**Figure 4:**
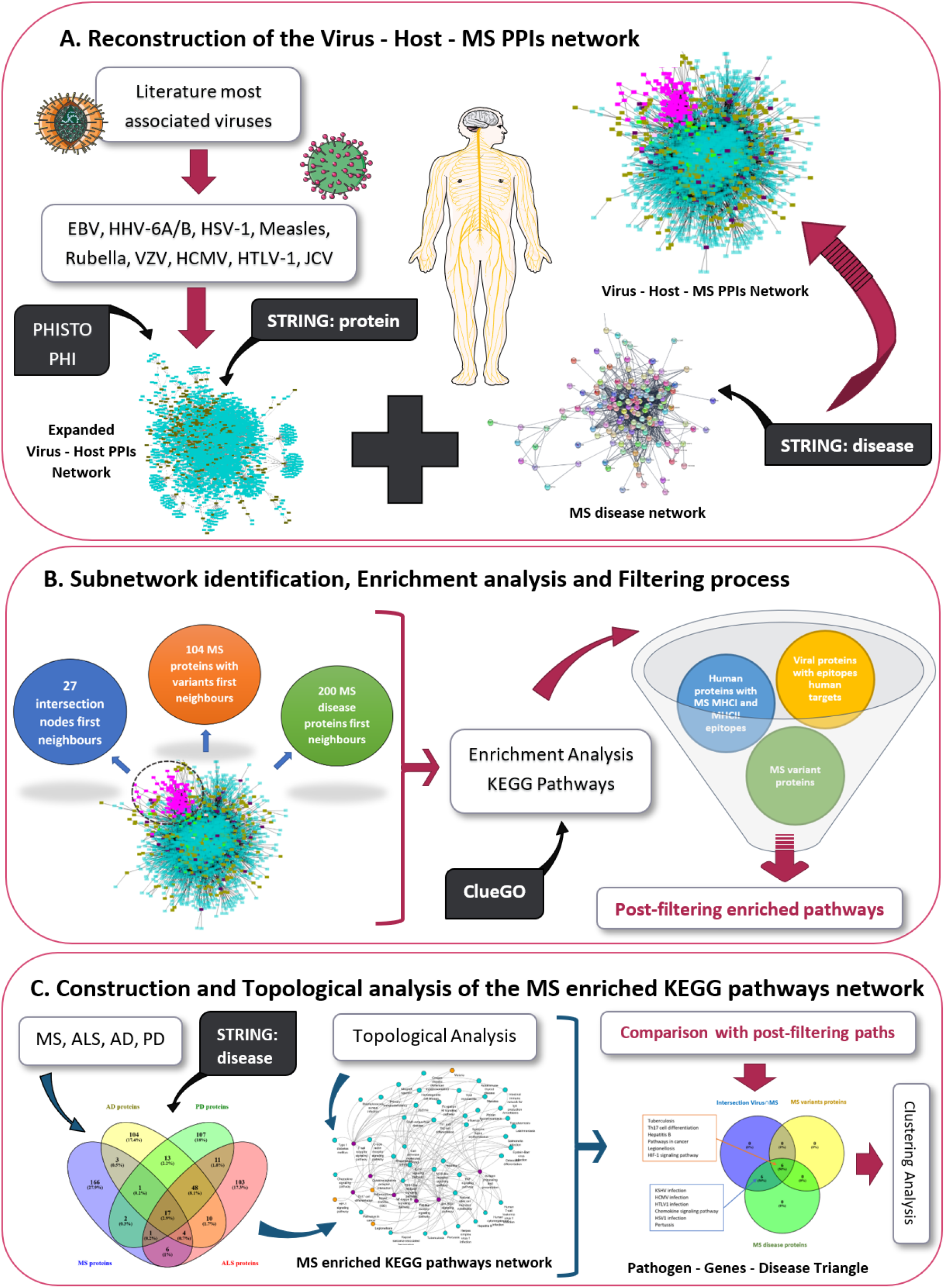
Schematic representation of the methodology applied in this paper to investigate the interaction between virus-host-MS PPIs using a network-based approach with the aim to identify viral-mediated pathogenic mechanisms that might be involved in the development of MS. Figure contains illustrations obtained from Servier medical art (https://smart.servier.com/), provided free and licensed under the Creative Commons Attribution 3.0 Unported License.

## METHODS

### Reconstruction and visualization of the integrated Virus - Host - MS PPIs network

A thorough literature review analysis has been performed to identify viruses highly associated with MS disease (Table 1). Eleven viruses have been identified. From them the Human Endogenous Retrovirus was excluded due to lack of PPI data. Virus-host PPI data for the ten selected viruses were collected from PHISTO database which is one of the most comprehensive databases for pathogen - human host PPI data and contains only experimentally detected interactions that are imported from several PPI databases [79,80]. For the selected viruses, a unified virus-host PPI network was constructed and visualized in Cytoscape [81]. The characteristics of the ten viruses and the number of virus-host PPIs for each of the viruses included are indicated in Table 2. The StringAPP [77] was then used to expand the interactome of the viral-targeted human proteins and create edges between the human proteins, in order to create an expanded interconnected virus-host PPI network. The unified virus-host PPIs network contained 2101 different viral-targeted human proteins, which were expanded by maximum of 10 additional interactors using the *STRING: protein* query and the confidence cut-off for the interactions between the human proteins was set at 0.8. The confidence cut-off score determines the nature and quality of the supporting evidence of the interaction between proteins and ranges from 0 (low) to 1.0 (high), with interactions with high confidence score being more likely to be true positives. Expansion of the 2101 viral-targeted human, resulted in the import of an additional 238 human proteins. Then the expanded interconnected virus-host PPIs network, containing 2339 human proteins and 295 viral proteins, was integrated with the MS disease PPI network that contained the 200 disease associated proteins with the highest disease scored obtained from the *STRING: disease* query of the StringApp. Merging of the two networks resulted in the construction of an integrated virus - host - MS PPI network, of 2807 nodes and 33741 edges.

**Table 2:** Viruses – Human host PPI network data

| <u>Virus species</u> | <u>Family</u> | <u>Virus genome</u> | <u># of strains</u> | <u># of PHIs</u> | <u># of viral proteins</u> | <u># of human proteins</u> |
| --- | --- | --- | --- | --- | --- | --- |
| Herpes Simplex virus 1 (HSV-1) | Herpesviridae | dsDNA | 6 | 777 | 66 | 604 |
| Varicella-zoster virus (HHV-3) | Herpesviridae | dsDNA | 2 | 6 | 5 | 4 |
| Epstein-Barr Virus (EBV) | Herpesviridae | dsDNA | 3 | 4644 | 148 | 1198 |
| Human Cytomegalovirus (HCMV) | Herpesviridae | dsDNA | 4 | 110 | 32 | 90 |
| Human Herpes Virus 6A (HHV-6A) | Herpesviridae | dsDNA | 2 | 3 | 3 | 3 |
| Human Herpes Virus 6B (HHV-6B) | Herpesviridae | dsDNA | 1 | 4 | 3 | 4 |
| John Cunningham virus (JCV) | Polyomaviridae | dsDNA | 1 | 51 | 5 | 49 |
| Rubella virus | Togaviridae | +ssRNA | 3 | 60 | 4 | 59 |
| Human T-cell lymphotropic virus type 1 (HTLV-1) | Retroviridae | +ssRNA-RT | 5 | 195 | 15 | 178 |
| Measles morbillivirus virus | Paramyxoviridae | -ssRNA | 6 | 529 | 14 | 487 |
| **Total** |  |  | **33** | **6379** | **295** | **2676** |

### Topological and knowledge - based subnetwork identification in the Virus - Host - MS PPI network

Due to the size and the complexity of the integrated virus-host-MS PPI network it is important to identify subnetworks that reveal important virus-host interactions that could directly or indirectly affect disease proteins. Three subnetworks were isolated that account for pathogen - genes - MS disease interactions:

1. The **first neighbors of the intersection nodes between the expanded virus-host PPI network and the MS disease PPI network:** merging of the expanded virus-host PPI network with the MS disease network indicated 27 intersection nodes, which are MS disease related proteins and 21 nodes are also human viral targets. This subnetwork, includes the 27 intersection nodes and their first neighbours.
2. The **first neighbors of MS related proteins**: the second subnetwork includes the 200 MS disease associated proteins and their first neighbours.
3. The **first neighbors of human proteins with MS related variants**: the MS related variants subnetwork was isolated by mapping on the integrated virus - host - MS PPI network, MS disease associated variant proteins, that were collected from the DisGeNET database [75]. This mapping revealed 104 MS variant proteins on the network which were then selected with their first neighbours to highlight the third subnetwork. Based on the local impact hypothesis, viral host targets of the associated virus are located in proximity of disease susceptibility genes within the network and the expression pattern of such genes changes significantly [37,46,82]. Therefore, viral proteins that directly interact with MS disease associated variant proteins or indirectly by affecting another protein within the same pathway of the disease variant can lead to a shift from virus-host equilibrium state to disease disequilibrium state, causing viral perturbations.

The number of viral and MS disease proteins included in each subnetwork can be found in Table 3.

**Table 3:**
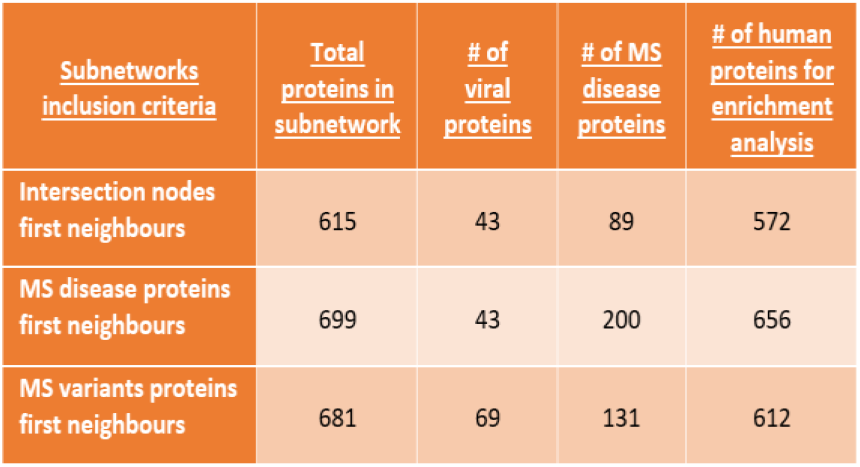
Contents of the topological and knowledge - based subnetworks.

| Subnetworks inclusion criteria | Total proteins in subnetwork | # of viral proteins | # of MS disease proteins | # of human proteins for enrichment analysis |
| --- | --- | --- | --- | --- |
| Intersection nodes first neighbours | 615 | 43 | 89 | 572 |
| MS disease proteins first neighbours | 699 | 43 | 200 | 656 |
| MS variants proteins first neighbours | 681 | 69 | 131 | 612 |

### Enrichment analysis of the three subnetworks

Pathway enrichment analysis was performed for the human proteins contained in each of the three subnetworks (Table 3) using the ClueGO plugin in Cytoscape [83]. Pathway enrichment analysis was performed using the KEGG pathways database [84] keeping only the terms with significant p −value ≤ 0.05 (corrected with Bonferroni step-down).

### Similarity and clustering analysis of viral proteins - KEGG pathways interactions

To identify clusters of viral proteins that interact with similar KEGG pathways, and vice versa, we used the vegan package in R [85] to measure the Jaccard similarity index. Then we performed agglomerative hierarchical clustering using the factoextra package in R [86].

### Reconstruction and visualization of the MS enriched pathway-to-pathway network

NDs share several common pathological mechanisms, therefore common disease proteins shared between all NDs could reflect direct commonalities of pathological mechanisms and pathways affected, whereas unique associated disease proteins might possibly reflect more specific disease phenotype. We selected AD (DOID: 10652), ALS (DOID:332), and PD (DOID:14330) to be compared to MS (DOID:2377) for the following reasons: (1) like MS, their development has also been associated with various viral infections, as indicated in Table 1, (2) similar to MS, they affect the CNS unlike other NDs that affect the peripheral nervous system, (3) they are also predominantly sporadic but also have familial forms unlike other NDs like Huntington’s Disease and Spinocerebellar Ataxia which are hereditary and (4) most importantly these 4 NDs share several common pathological mechanisms. The *STRING: disease* query of the StringApp in Cytoscape was used to collect disease associated proteins for the four NDs. The top 200 most associated disease proteins with the highest disease score for each disorder were collected, with confidence cut – off set at 0.8. We used Venny 2.1 [87], to compare the associated proteins among the four NDs to identify the common disease associated proteins between all four NDs and those that are unique for MS compared to this specific group of NDs (Figure 5).

**Figure 5:**
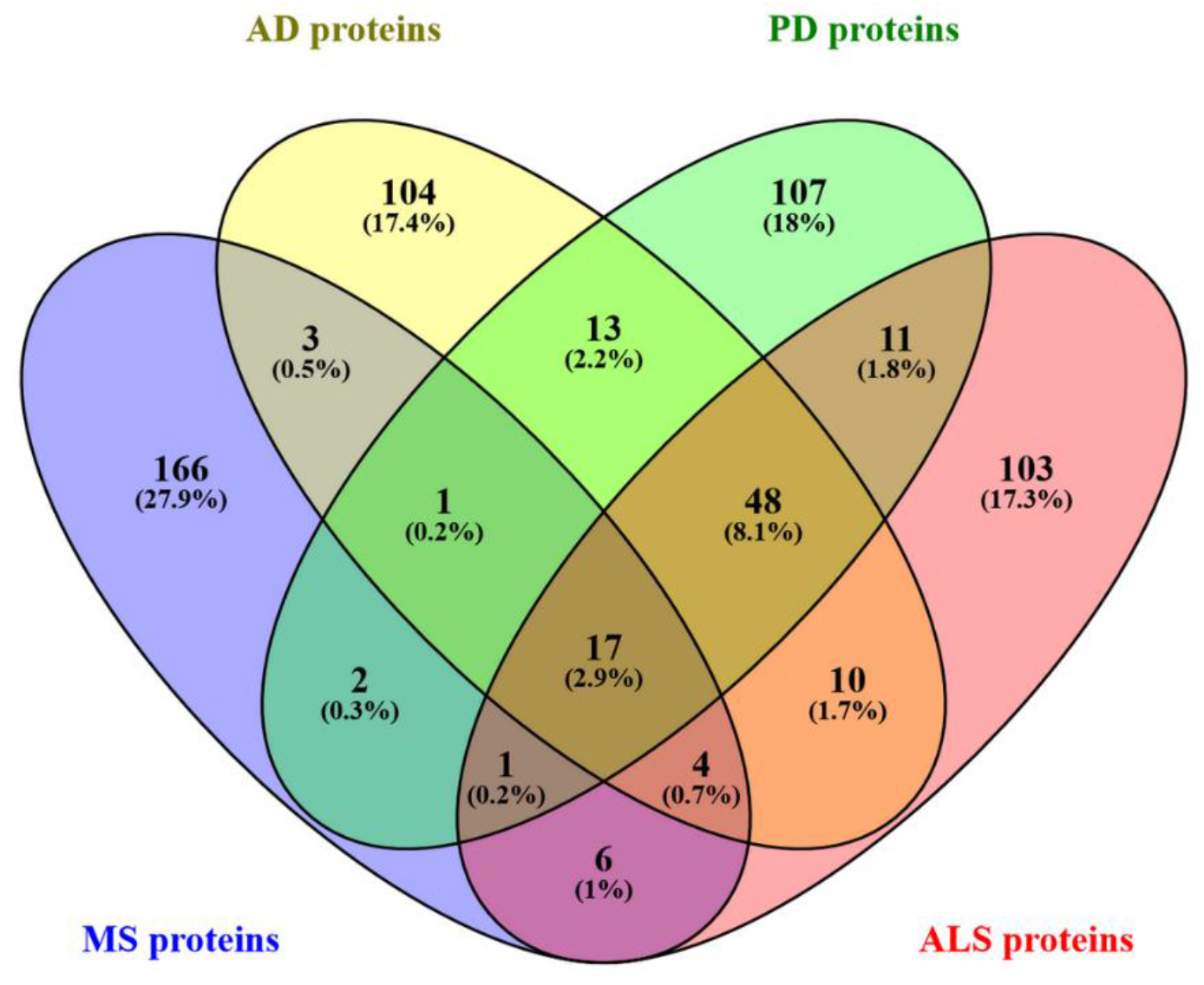
Comparison results of the top 200 highest associated disease proteins between the four NDs: ALS, MS, PD, and AD.

We identified 17 common disease proteins between all four NDs proteins, 166 unique to MS and 17 that are shared between MS and some of the other NDs. Enrichment analysis was performed for each group of proteins with the ClueGO app [83] in Cytoscape using the KEGG pathways database [84] keeping only the terms with significant p −value ≤ 0.05 (corrected with Bonferroni step-down) (Figure 6). The enriched KEGG pathway results obtained from the 17 common ND proteins were then entered into the PathwayConnector web tool [88] to discover a complementary network of pathways that interact with the common ND pathways (Figure 7).

**Figure 6:**
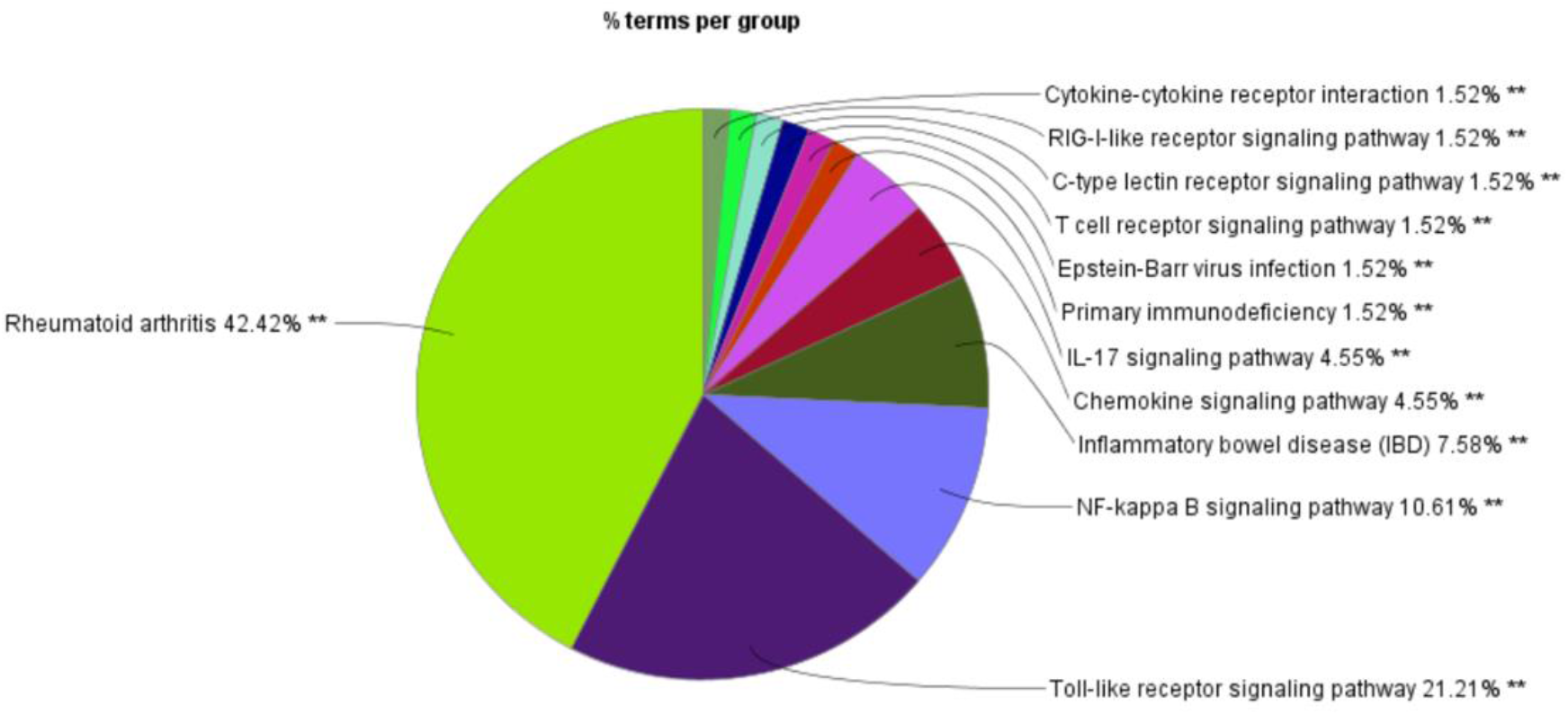
Enriched KEGG pathways analysis results of the 166 unique MS disease proteins, obtained using the ClueGO app in Cystoscope, with the pathways classified into groups and the percentage indicating the number of terms in each group.

**Figure 7:**
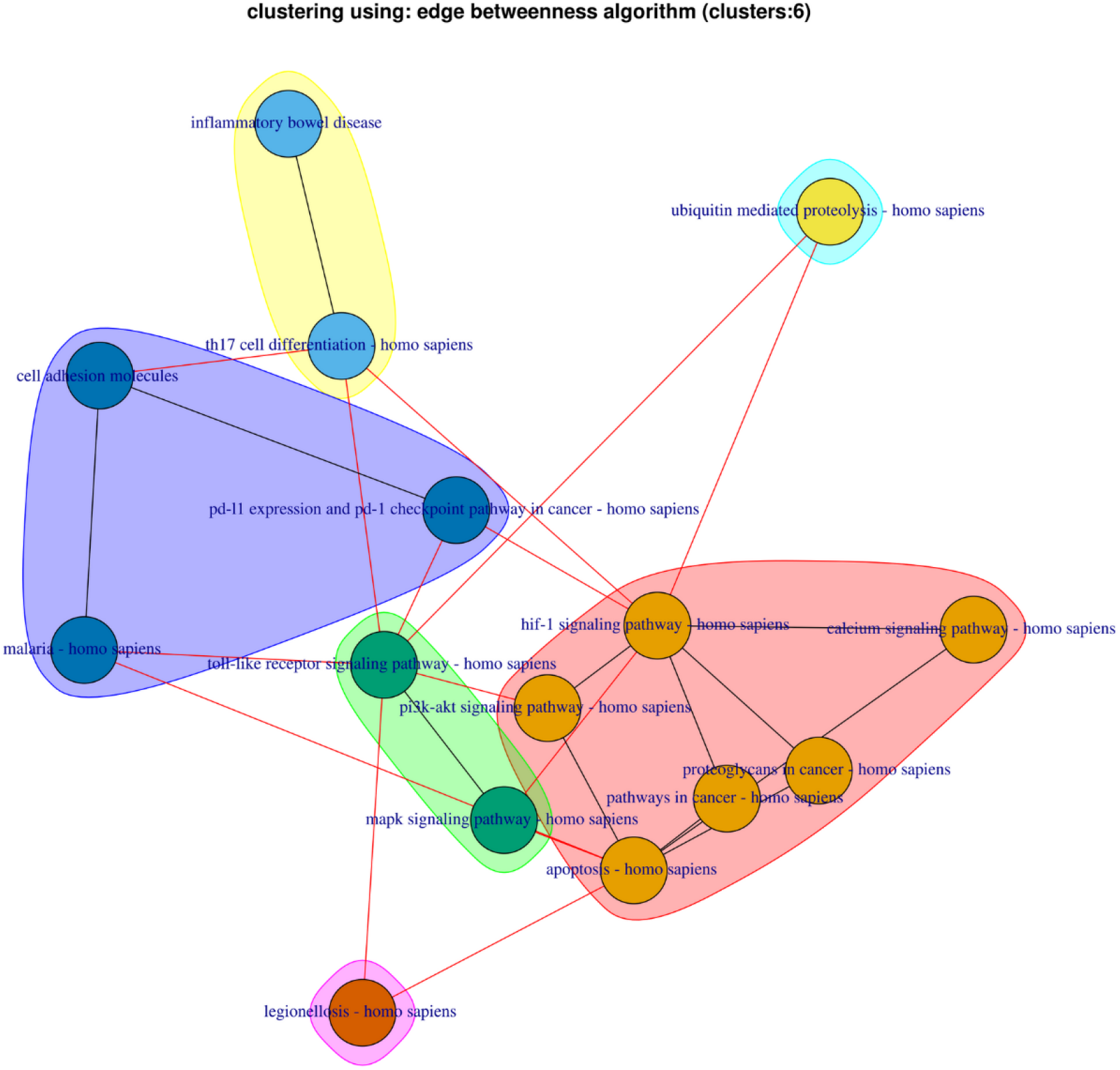
Complementary network of the 4 common NDs KEGG pathways (IBD, HIF-1 signaling pathway, Malaria and Legionellosis) and their interactions with the 11 complementary nodes/pathways, created using PathwayConnector.

Moreover, PathwayConnector was used to reconstruct and visualize the MS enriched KEGG pathways network. Topological analysis was performed using the CytoHubba app of Cytoscape [89] with the aim to identify hubs and bottlenecks nodes in the MS enriched KEGG pathways network. The ClusterMaker app [90] was then used to perform community clustering [91] on the MS enriched KEGG pathways network, which detects clusters of nodes based on their connectivity, allowing to identify community clusters of pathways that interact with the hub-bottleneck nodes.

## RESULTS

### Enrichment analysis results of the MS unique, MS shared and NDs common disease proteins

The enrichment analysis of the 17 common NDs associated proteins resulted in the identification of 4 common KEGG pathways: Inflammatory Bowel Disease (IBD), Malaria, HIF-1 signaling pathway and Legionellosis pathways. Complementary network reconstruction of the 4 KEGG pathways indicated that they interact with 11 complementary pathways (Figure 7). Enrichment analysis of the 166 MS unique disease associated proteins revealed 49 KEGG pathways (Figure 6) which are either infectious disease or immune related pathways (see supplementary data), thus confirming the association between pathogens and their interaction with the immune system in the development of MS. Interestingly, 42.4% of KEGG pathways enriched results belong to the Rheumatoid arthritis group. In addition, the enrichment analysis results of the 17 disease associated proteins that are shared between MS and some of the other NDs resulted in 7 KEGG pathways.

### Topological and clustering analysis of the MS enriched KEGG pathways network

Reconstruction and topological analysis of the MS enriched KEGG pathways network allowed to identify the most important bottleneck and hub pathways within the network. Topological analysis revealed 7 hub-bottleneck nodes/pathways that can act as a bridge of communication between the rest of the pathways in the network. This high centrality nodes were identified by using the average score of the top 10 Bottleneck, Degree and Closeness centrality pathway results obtained via the CytoHubba app [89]. We also examined the topological characteristics of the 4 common NDs pathways in the MS enriched KEGG pathways network which indicated that Malaria, HIF-1 signaling pathway and Legionellosis pathways are non-hub-non-bottleneck nodes, whereas the IBD pathway acts as a hub-non-bottleneck node.

Clustering analysis of the MS enriched KEGG pathways network using the community clustering algorithm [91] allowed to identify 4 clusters of community nodes that interact with the 7 hub-bottleneck nodes. The clustering results indicate that the majority of viral infectious disease pathways are clustered together with the hub-bottleneck nodes Toll-like receptor signaling pathway and JAK-STAT signaling pathways (Figure 8a), whereas few viral infectious disease pathways are located in cluster c, with the hub-bottleneck nodes Th17 cell differentiation and NF-kappa B signaling pathway (Figure 8c).

**Figure 8:**
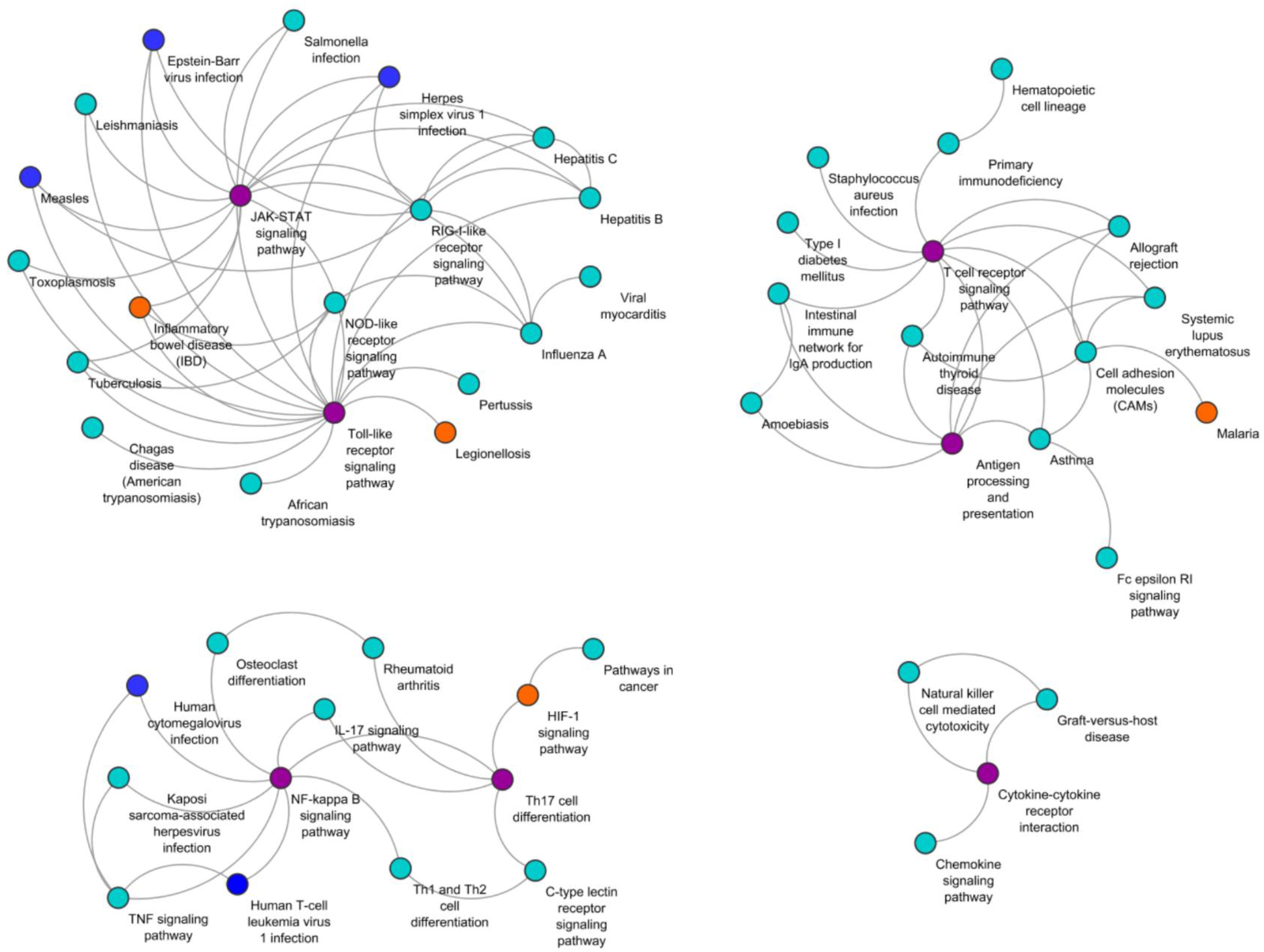
Community clustering of the MS enriched KEGG pathways network resulted in the formation of 4 community clusters. Orange color nodes represent the 4 common NDs pathways between ALS, MS, PD, AD. Hub-bottleneck nodes are represented in color purple, and blue color nodes represent infectious disease pathways of some of the viruses we included for the reconstruction of our virus-host-MS PPIs network.

### Filtering process of the topological and knowledge-based subnetworks enrichment results

The enrichment analysis of the human proteins contained in each subnetwork (Table 3) revealed multiple significant KEGG pathways. A filtering process (Figure 9) was performed on the enriched terms for each of the three subnetworks in an effort to isolate the most relevant KEGG pathways. The selection criteria are listed below:

1. **Criterion 1:** only KEGG pathways containing human proteins that are targets of immunogenic viral proteins were selected. To identify the immunogenic viral proteins, immune epitope data for the selected viruses that are found in humans during viral infection where obtained from the Immune Epitope Database (IEDB) [92].
2. **Criterion 2:** The results obtained from the first filtering process where then filtered by using immune epitope data found in humans with MS disease. Immune epitope data found in MS disease individuals, that have MHCI or MHCII - restricted antigen recognition, meaning that T cells will only respond to the antigens from this protein only when they are bound to the relevant MHC molecules, were obtained from the IEDB [92]. The epitope data contained few epitopes against viral proteins where their human targets were selected as in criterion 1, but the majority were self-epitopes against human proteins which were also selected.
3. **Criterion 3:** Finally, the filtering results obtained were filtered to select pathways containing MS variants proteins collected previously from the DisGeNET database [75].

**Figure 9:**
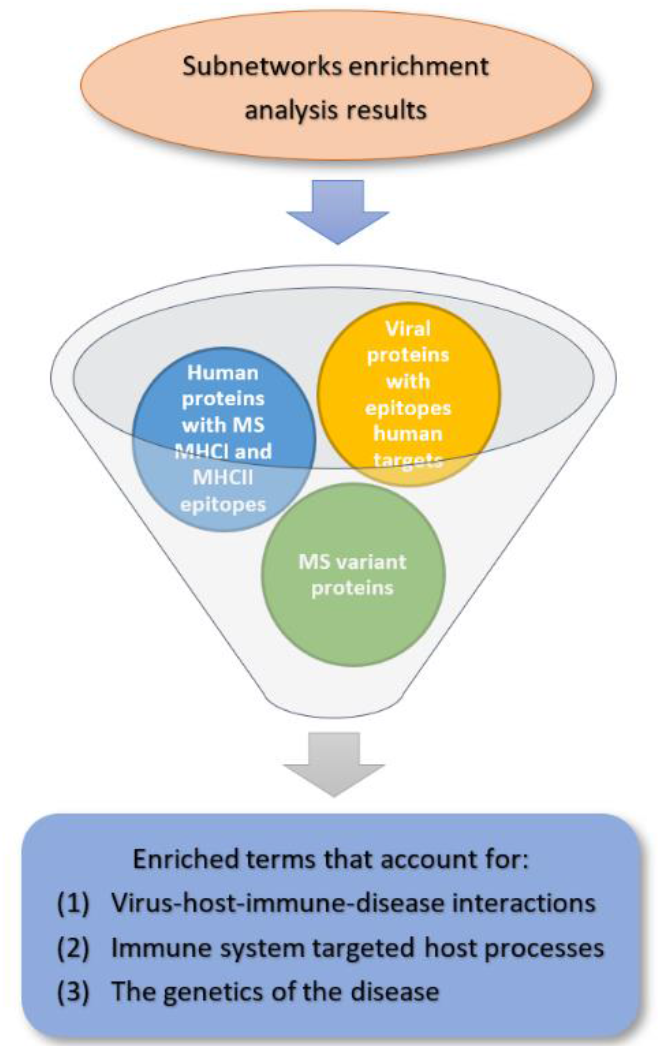
Filtering process applied on the enriched KEGG pathways analysis results of the three subnetworks.

This 3-criteria filtering process was applied on the intersection subnetwork and the MS disease proteins subnetwork, whereas for the MS variant proteins subnetwork only the first 2-criteria where applied. The filtering process aims to account for the immunogenicity, autoimmunity and genetic components of MS disease as the final pathways obtained contain immunogenic viral proteins and their human targets, host proteins with self-epitopes and MS disease variants proteins.

### Identification of post-filtering enrichment results that are also MS enriched KEGG pathways

The filtering process significantly reduced the number of enriched KEGG pathways for the three subnetworks, allowing possibly to isolate the most relevant pathways. More specifically, the enriched KEGG pathways before the filtering process for the intersection subnetwork that contains nodes that are both human viral targets and MS-related proteins were 94, for the MS disease proteins and MS variants proteins subnetworks 91 and 94, whereas after the filtering process 36, 34, 28 KEGG pathways remained, respectively. Comparison of the post-filtering enrichment results for each of the three subnetworks with the MS unique, MS shared and common NDs pathways that were previously found, allowed to identify and select enriched terms from the three subnetworks that are also MS disease related terms.

### Comparison of the enriched pathways between the three subnetworks

The three subnetworks represent the triangle to pathogenesis by accounting the importance of three elements, pathogen - genes - disease interactions, and their role in the pathogenesis of MS disease (Figure 10a). To identify the KEGG pathways that fall within the pathogen - genes - disease triangle, we compared the post-filtering enrichment results that are also MS disease related terms between the three subnetworks, which resulted in the identification of 12 KEGG pathways (Figure 10b).

**Figure 10:**
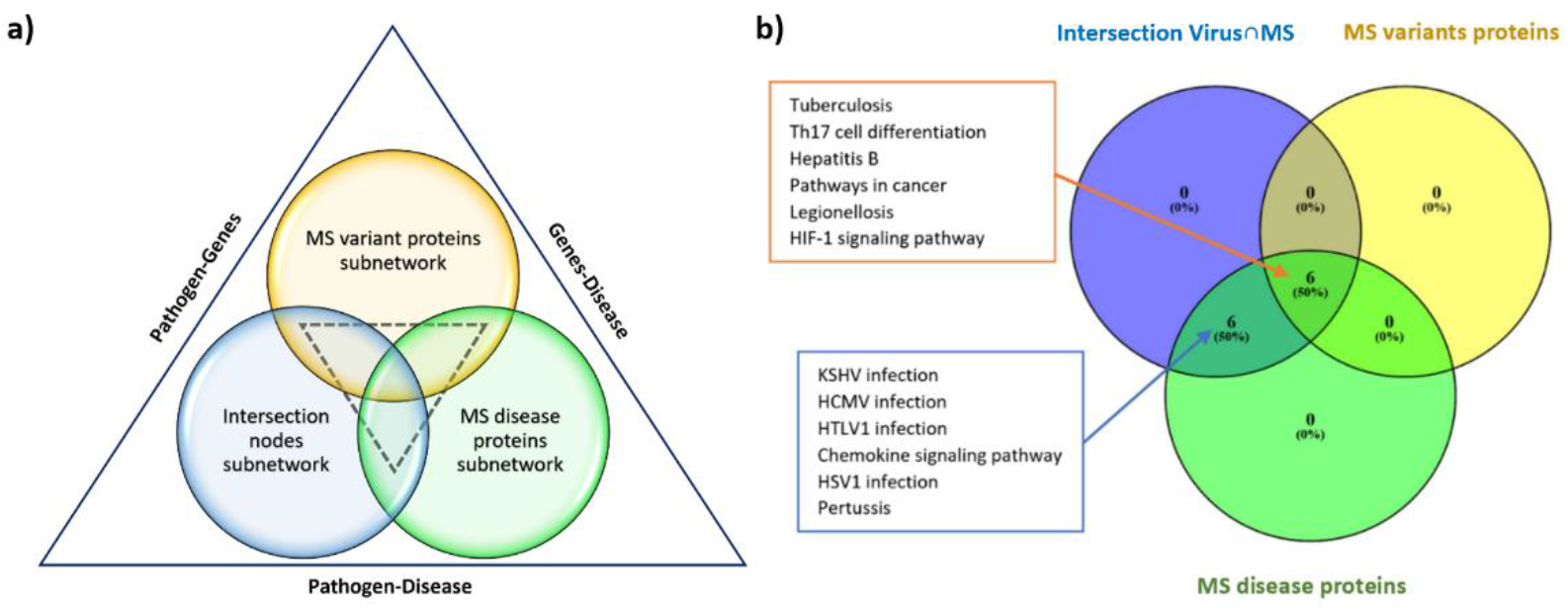
(a) Pathogen - Genes - Disease triangle. (b) Comparison of the post-filtering KEGG pathways enriched results that are also MS disease related terms between the three subnetworks, indicating the 12 KEGG pathways that fall within the Pathogen - Gene - Disease triangle.

### Analysis of the complementary network of the final 12 KEGG pathways

Pathway-to-pathway network reconstruction of the 12 KEGG pathways using the PathwayConnector tool indicated that only Th17 cell differentiation pathway, HIF-1 signaling pathway and Pathways in cancer functionally interact, whereas the rest 9 KEGG pathways are disconnected with each other. A complementary network was therefore created by using the missing pathway approach to identify proximal pathways that interact with the 12 KEGG pathways, which led to the identification of 7 complementary pathways, shown in Table 4. Comparison of the complementary pathways with the MS disease related pathways, showed that 4 out of the 7 pathways are also MS unique pathways, whereas 3 of the complementary pathways, namely TGF-beta signaling, Calcium signaling and MAPK signaling pathway do not belong to the MS disease pathway terms (Table 4).

**Table 4:** Comparison of the 7 complementary pathways (*) identified in the 12 KEGG pathways complementary network with the MS disease pathways terms. Combined p-value obtained from the three subnetworks and whole network p-values of the 12 KEGG pathways and the 7 complementary pathways (*).

| KEGG pathway name | MS unique, MS shared, NDs common | Average p-value | Whole network p-value <0.05 | Whole network p-value >0.05 |
| --- | --- | --- | --- | --- |
| Tuberculosis | MS unique | 4.05E-14 | 5.29E-05 | - |
| KSHV infection | MS unique | 2.87E-17 | 9.03E-08 | - |
| Th17 cell differentiation | MS unique | 3.00E-14 | 4.08E-05 | - |
| HCMV infection | MS unique | 6.57E-16 | 7.79E-10 | - |
| Hepatitis B | MS unique | 6.02E-22 | 1.27E-13 | - |
| HTLV-1 infection | MS unique | 1.65E-15 | 3.98E-17 | - |
| Chemokine signaling pathway | MS unique | 1.02E-09 | - | 8.09E-01 |
| HSV-1 infection | MS unique | 4.42E-27 | 5.68E-28 | - |
| Pathways in cancer | MS unique | 9.92E-14 | 3.16E-05 | - |
| Pertussis | MS shared | 1.40E-06 | - | 6.86E-01 |
| Legionellosis | MS unique, MS shared, NDs common | 6.31E-11 | 7.15E-08 | - |
| HIF-1 signaling pathway | NDs common | 1.03E-04 | 3.05E-02 | - |
| TGF-beta signaling pathway* | - | 1.01E-02 | - | 6.05E-02 |
| T-cell receptor signaling pathway* | MS unique | 4.71E-12 | 7.08E-04 | - |
| Calcium signaling pathway* | - | - | - | 1.00E+00 |
| JAK-STAT signaling pathway* | MS unique | 8.74E-10 | - | 3.90E-01 |
| MAPK signaling pathway* | - | 1.97E-05 | - | 2.10E-01 |
| NF-kappa B signaling pathway* | MS unique | 5.47E-17 | 1.32E-08 | - |
| Toll-like receptor signaling pathway* | MS unique | 8.53E-19 | 6.67E-09 | - |

Topological analysis of the MS enriched KEGG pathways network, indicated 7 hub-bottleneck nodes that act as a bridge of communication within the network, 5 of these hub-bottleneck nodes are found in the complementary network, of which 4 are complementary nodes and 1 is part of the 12 KEGG pathways. A schematic visualization (Figure 11) illustrates the complementary network of the 12 KEGG pathways.

**Figure 11:**
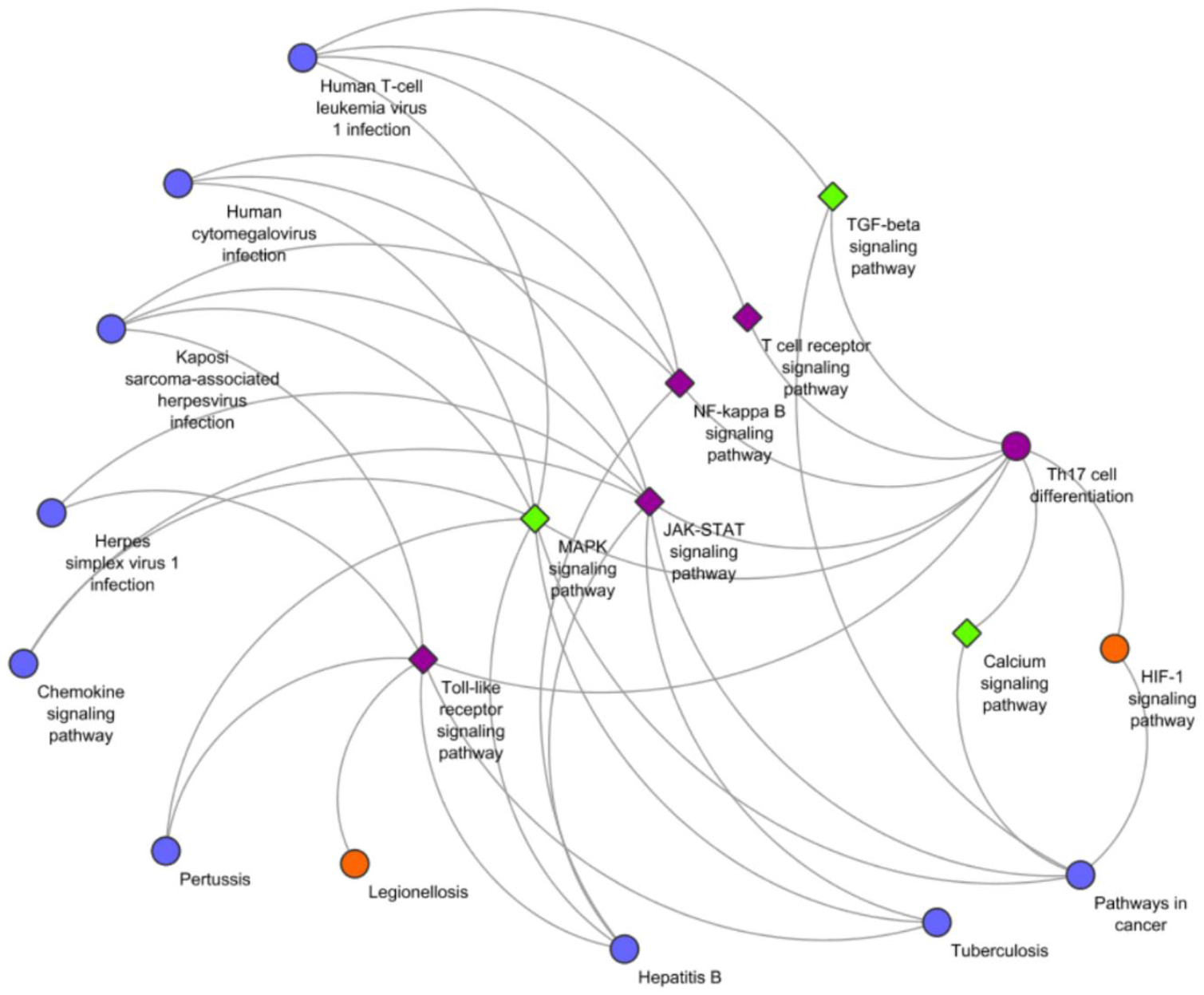
Schematic visualization of the complementary network for the 12 KEGG pathways, which are indicated in circular nodes, and the 7 complementary nodes shown in diamond shape. The orange nodes represent nodes that are also part of the 4 common NDs pathways and purple nodes are hub-bottleneck nodes that act as a bridge of communication between nodes/pathways in the MS disease KEGG pathways network. The green nodes are complementary nodes which are not MS disease related pathway terms (MS unique, MS shared or NDs common).

### Similarity and clustering results of viral proteins-KEGG pathways interactions

The final 12 KEGG pathways are targeted by 67 viral proteins from 8 viral species, more specifically EBV, HCMV, HHV-6A, HHV-6B, HSV-1, HTLV-1, Measles and Rubella, with the majority being EBV strains. The final step of our analysis involved identifying clusters within the 67 viral proteins that target similar KEGG pathways and clusters within the 12 KEGG pathways that are targeted by similar viral proteins. Similarity analysis was performed using the Jaccard similarity index and hierarchical clustering analysis for both dendrograms was preformed using the average distance (also called mean) method as it had the highest correlation coefficient value in both cases. The clustering dendrogram in Figure 12 indicates the presence of 16 clusters of viral proteins-based on pathway target similarity with the 12 KEGG pathways. The clustering dendrogram of the 12 KEGG pathways and a heatmap plot indicating the interactions between the 67 viral proteins with the 12 KEGG pathways can be found in the supplementary data.

**Figure 12:**
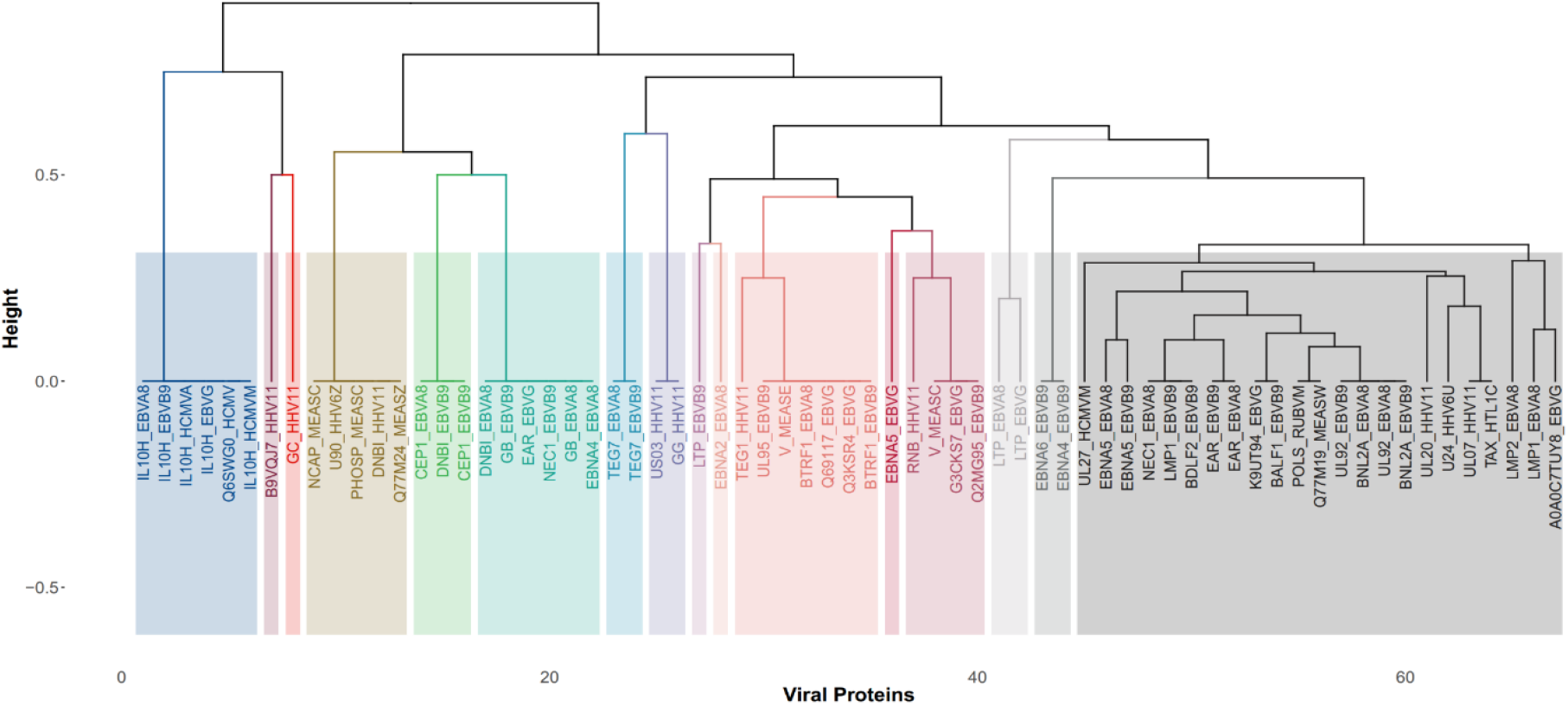
Clustering dendrogram of the 67 viral proteins from 8 viral species EBV, HCMV, HHV-6A, HHV-6B, HSV-1, HTLV-1, Measles and Rubella based on target similarity of the final 12 KEGG pathways.

## Discussion

In this study, we presented an integrative network-based bioinformatics pipeline approach that utilizes various databases and bioinformatics tools presented in the first part of this paper with the aim to identify viral - mediated pathogenic mechanisms that may be associated with the development and/ or progression of MS disease. To investigate viral-mediated perturbation in MS disease we compared the disease associated proteins between four NDs; ALS, MS, PD and AD that have been associated with viral infections in order to identify MS unique, MS shared and NDs common pathways. We identified 4 common ND KEGG pathways that may represent common pathological mechanisms: the IBD, HIF-1 signaling pathway, Malaria and Legionellosis pathways. Interestingly, all 4 NDs have been associated with IBD which is characterized by chronic inflammation in the gastrointestinal gut and one main hypothesis is that abnormal brain-gut interactions might be involved in its pathogenesis [93–97]. The HIF-1 signaling pathway on the other hand is involved in the regulation of molecular signals during hypoxia, infection and the stimulation of pro-inflammatory signals [98]. Hypoxia and the HIF-1 signaling pathway have been associated with the development and progression of wide range of diseases, with hypoxia being a common characteristic in NDs [99]. Malaria and Legionellosis are infectious pathways, thus highlighting the possible association between pathogens and NDs.

Statistically significant enrichment analysis of the 166 unique MS disease proteins also confirmed the association between pathogens and their interaction with the immune system in the development of MS. In addition, the KEGG enrichment results indicates that 42.4% of the pathways belong to the Rheumatoid arthritis group, suggesting the existence of common pathogenic mechanisms between MS and Rheumatoid arthritis [100–102].

Topological analysis of the MS enriched KEGG pathways network allowed to identify 7 hub-bottleneck nodes (Figure 13) that can act as a bridge of communication between the rest of the pathways in the network. Pathogens are known to target hubs and bottlenecks proteins in order to exert systemic infectious effects [44,49,55], therefore hub-bottleneck nodes can possibly act as disease communicator nodes bridging the communication between infectious pathways and the rest of the MS enriched KEGG network pathways. Subsequently, any dysregulation in a pathway that interacts with a hub-bottleneck or dysregulation of a hub-bottleneck node directly can lead to a cascade of events causing systemic dysregulation of a subset of pathways, resulting possibly in the development of MS. In order to identify communities of infectious disease pathways that interact with the identified hub-bottleneck pathways, we performed community clustering analysis of the MS enriched KEGG pathways network that indicated 4 clusters (Figure 8). More specifically, the community clustering results showed that EBV, HSV-1 and Measles viral infection pathways which have being linked as a risk factor for the development of MS interact with the Toll-like receptor signaling pathway and JAK-STAT signaling pathway hub-bottleneck nodes (Figure 8a). Whereas, HCMV and HTLV-1 that are also associated with MS form a different community cluster and interact with the NF-kappa B signaling pathway and Th17 cell differentiation pathway hub-bottleneck nodes (Figure 8c).

**Figure 13:**
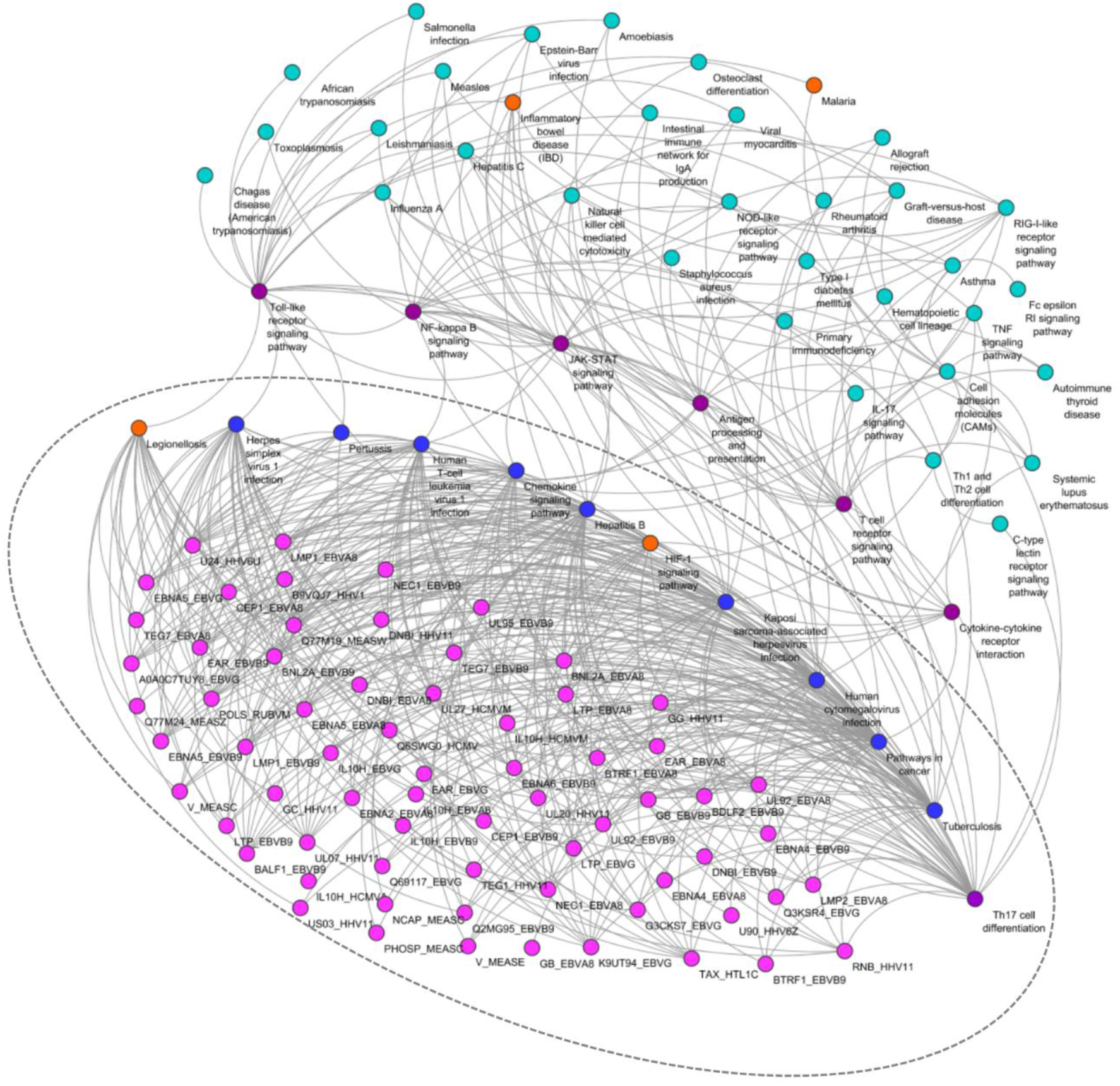
Schematic illustration of the possible viral-mediated pathogenic mechanisms obtained through our pipeline approach indicating how the resulting 67 viral proteins (lila colour) targeting the 12 identified KEGG pathways in the MS enriched KEGG pathways network which in turn interact with the hub - bottleneck disease communicator nodes (purple colour) can exert systemic effects within the network and lead to the development of MS. The NDs common pathways are indicated in orange.

Our methodology also involved the reconstruction, subnetwork identification and pathway enrichment analysis of the integrated virus-host-MS PPI network, where after applying our filtering process and isolating pathways that are also MS disease enriched terms, we were able to identify 12 enriched KEGG pathways (Figure 10b). The final KEGG pathways list obtained through our pipeline approach accounts for pathogen - genes - disease interactions as they are pathways that contain MS variants proteins, MS disease proteins and virus-host PPIs. Furthermore, via our filtering process, the final results account for the immunogenicity, autoimmunity and genetic components of MS disease. Finally, via the filtering process and the comparison with other NDs by obtaining only pathways that are associated with MS unique, MS shared and NDs common pathways, we were able to isolate the most relevant MS disease pathways from a large pool of enriched pathway results.

To investigate pathway-to-pathway interactions of the final 12 KEGG pathways, a complementary network was created as it can provide possible pathways of communication of the 12 KEGG pathways with other pathways, particularly MS disease related pathways and hub-bottleneck disease communicator nodes. The complementary network, shown in Figure 11, indicates 7 proximal complementary nodes with JAK-STAT signaling pathway, Toll-like receptor signaling pathway, NF-kappa B signaling pathway and T-cell receptor signaling complementary nodes being also MS disease related pathways and hub-bottleneck disease communicator nodes in the MS enriched KEGG pathways network. All the 12 KEGG pathways directly interact with one or more of the hub-bottleneck disease communicator nodes, therefore by interacting with disease communicator nodes they can exert systemic effects within the MS enriched KEGG pathways network (Figure 11). In addition, the three complementary nodes, Calcium signaling pathway, MAPK signaling pathway and TGF-β signaling pathway that are not MS disease related pathways interact with the Th17 cell differentiation pathway which is part of the 12 final KEGG pathways and a disease communicator node in the MS enriched KEGG pathways network, therefore they might indirectly contribute to the development of MS (Figure 11). The Th17 cell differentiation pathway acts as a key disease communicator node in the complementary network of the final 12 KEGG pathway as it interacts with all 7 of the complementary nodes of which 4 are also disease communicator nodes. Moreover, it interacts with the HIF-1 signaling pathway one of the four common NDs pathways, thus making it a central player as a possible viral-mediated pathogenic pathway in MS. The relapsing-remitting MS phase of MS involves the infiltration of CD4+ T cells with Th1 and Th17 proinflammatory phenotype [103,104], thus highlighting the role of the Th17 cell differentiation pathway in the immunopathogenesis of MS. Th17 cells are also involved in the immunopathogenesis of multiple other autoimmune diseases including IBD and Rheumatoid arthritis [105]. In addition, interferon beta which is used as a first line of treatment for MS patients at the relapsing-remitting MS phenotype have shown to exert its pharmacological effects by inhibiting indirectly Th17 cell differentiation [106–108].

The final 12 KEGG pathways are targeted by 67 viral proteins from 8 viral species. Therefore, viruses by targeting the identified pathways in the MS enriched KEGG pathways network, illustrated in Figure 13, which in turn interact with the hub-bottleneck disease communicator pathways may be able to exert systemic pathogenic effects, thus leading to the development of MS. By performing similarity and clustering analysis we were able to identify the group of KEGG pathways that can be targeted by each viral protein, but also clusters of viral proteins from different species that can target similar pathways. The clustering analysis also indicates viral proteins that can possibly exert similar effects in the MS disease pathways network by affecting the same group of pathways.

## Pipeline Limitations

One important limitation of our study is the imbalance of available experimental virus-host PPIs data, as more studied viruses like EBV have more available data, which accounts for 72.8% of the virus-host PPI data included in our integrated virus-host-MS PPI network, than less studied viruses, such as HHV-6A and HHV6B. A current approach used to overcome this limitation is to use computationally predicted virus-host PPIs obtained from machine learning algorithms, but such methods are not without drawbacks due to the high false positive and false negative rates [109]. Another limitation inherent to pathway enrichment analysis is that the standard approach is to select only statistically significant pathways with p-value ≤ 0.05. However, pathways that are not revealed through the current analysis as statistically significant could also be important as viral proteins also interact with human protein targets in these pathways and therefore might be able to cause disease effects via non-statistically significant pathways.

## Conclusion

In this paper we initially reviewed databases and tools that can be utilized to investigate viral-mediated perturbations of the host’s interactome that lead to the generation of complex diseases, such as NDs. We then presented our integrative network-based bioinformatics pipeline approach that accounts for pathogen - genes - disease PPIs with the aim to identify possible viral-mediated pathogenic mechanisms in MS disease.

Comparison between disease associated proteins of four NDs (ALS, MS, PD and AD) associated with viruses confirmed the role of pathogens in the development of NDs and led to the identification of the HIF-1 signaling pathway as a possible common pathogenic mechanism in NDs. Reconstruction and topological analysis of the MS enriched KEGG pathways network that includes MS unique, MS shared and NDs common pathways enabled us to identify 7 hub-bottleneck nodes that can act as disease communicator nodes and exert systemic effects.

Through the reconstruction of the virus - host - MS PPIs network and the application of our methodology we were also able to identify 12 KEGG pathways targeted by 67 viral proteins from 8 viral species. These viruses might exert their viral-mediated pathogenic mechanisms by interacting with these 12 KEGG pathways which in turn interact with the hub-bottleneck disease communicator nodes, allowing them to exert systemic effects, hence affecting several MS disease pathways.

Finally, our analysis highlights the Th17 differentiation pathway which is part of the 12 underlined KEGG pathways as a key viral-mediated pathogenic mechanism and a possible therapeutic target for MS disease as it is a central hub-bottleneck disease communicator node in the MS disease enriched KEGG pathways network. To further elucidate the role of these pathways in MS disease, computational modelling can be used to determine if the viral proteins that target a specific pathway can shift the pathway into a disequilibrium state causing viral-mediated perturbations in the host interactome.

## Key points

- NDs are chronic degenerative neurological diseases and currently there are no effective pharmacotherapies for their treatment.
- Although several environmental and genetic factors have been implicated to their development, the exact underlying mechanisms are still unclear.
- Viral infections have been associated with several NDs, with virus-host PPIs representing a key pathogenic mechanism that can lead to the generation of perturbations within the human interactome.
- This work illustrates how network-based approaches used for the investigation of NDs and virus-host PPIs can be integrated to identify viral-mediated pathogenic mechanisms in MS.

## Data Availability Statement

Data available on request.

## Supplementary Data

Supplementary data attached in separate file.

