## Supplementary material for "Identification of viral-mediated pathogenic mechanisms in neurodegenerative diseases using network-based approaches": Suplemmentary data

### Supplementary data

**Table 1**: Enriched KEGG pathway terms and p-values for the 166 unique MS disease proteins and categorization of each pathway

| KEGG pathway term | *P*-value | Term category |
|---|---|---|
| Cytokine-cytokine receptor interaction | 5.72E-25 | Immune system |
| RIG-I-like receptor signaling pathway | 0.0153666 | Immune system |
| C-type lectin receptor signaling pathway | 0.0226666 | Immune system |
| T cell receptor signaling pathway | 1.68E-08 | Immune system |
| Epstein-Barr virus infection | 4.30E-07 | Infectious disease |
| Primary immunodeficiency | 4.29E-05 | Immune system disease |
| IL-17 signaling pathway | 5.71E-09 | Immune system |
| Fc epsilon RI signaling pathway | 0.0137505 | Immune system |
| TNF signaling pathway | 5.26E-08 | Immune system |
| Chemokine signaling pathway | 1.86E-07 | Immune system |
| Chagas disease (American trypanosomiasis) | 1.95E-06 | Infectious disease |
| Human cytomegalovirus infection | 0.0010681 | Infectious disease |
| JAK-STAT signaling pathway | 2.01E-09 | Immune system role |
| Th1 and Th2 cell differentiation | 5.00E-09 | Immune system |
| Measles | 6.88E-09 | Infectious disease |
| Pathways in cancer | 4.43E-05 | Cancer disease |
| Inflammatory bowel disease (IBD) | 1.15E-19 | Autoimmune Disease |
| NF-kappa B signaling pathway | 7.38E-09 | Immune system role |
| NOD-like receptor signaling pathway | 5.40E-04 | Immune system |
| Salmonella infection | 2.05E-04 | Infectious disease |
| Legionellosis | 0.0044721 | Infectious disease |
| African trypanosomiasis | 4.29E-05 | Infectious disease |
| Malaria | 1.83E-06 | Infectious disease |
| Osteoclast differentiation | 0.0038966 | Immune system activated |
| Toll-like receptor signaling pathway | 1.65E-16 | Immune system |
| Natural killer cell mediated cytotoxicity | 0.0044706 | Immune system |
| Leishmaniasis | 5.99E-07 | Infectious disease |
| Tuberculosis | 1.31E-12 | Infectious disease |
| Hepatitis C | 7.40E-04 | Infectious disease |
| Hepatitis B | 0.0011036 | Infectious disease |
| Influenza A | 5.62E-10 | Infectious disease |
| Kaposi sarcoma-associated herpesvirus infection | 5.52E-06 | Infectious disease |
| Herpes simplex virus 1 infection | 1.26E-07 | Infectious disease |
| Cell adhesion molecules (CAMs) | 1.69E-13 | Immune system |
| Antigen processing and presentation | 0.0231460 | Immune system |
| Hematopoietic cell lineage | 1.11E-07 | Immune system role |
| Th17 cell differentiation | 1.42E-12 | Immune system |
| Intestinal immune network for IgA production | 1.72E-14 | Immune system |
| Type I diabetes mellitus | 5.97E-07 | Autoimmune Disease |
| Toxoplasmosis | 6.35E-07 | Infectious disease |
| Staphylococcus aureus infection | 2.77E-04 | Infectious disease |
| Human T-cell leukemia virus 1 infection | 8.64E-04 | Infectious disease |
| Asthma | 6.40E-07 | Autoimmune Disease |
| Autoimmune thyroid disease | 9.23E-10 | Autoimmune Disease |
| Systemic lupus erythematosus | 2.98E-05 | Autoimmune Disease |
| Rheumatoid arthritis | 3.93E-15 | Autoimmune Disease |
| Allograft rejection | 3.42E-13 | Immune system mediated |
| Graft-versus-host disease | 6.02E-06 | Immune system mediated |
| Viral myocarditis | 8.90E-06 | Infectious disease |

**Figure 1:** Clustering dendrogram of the final 12 KEGG pathways based on interactions similarity with the 67 viral proteins

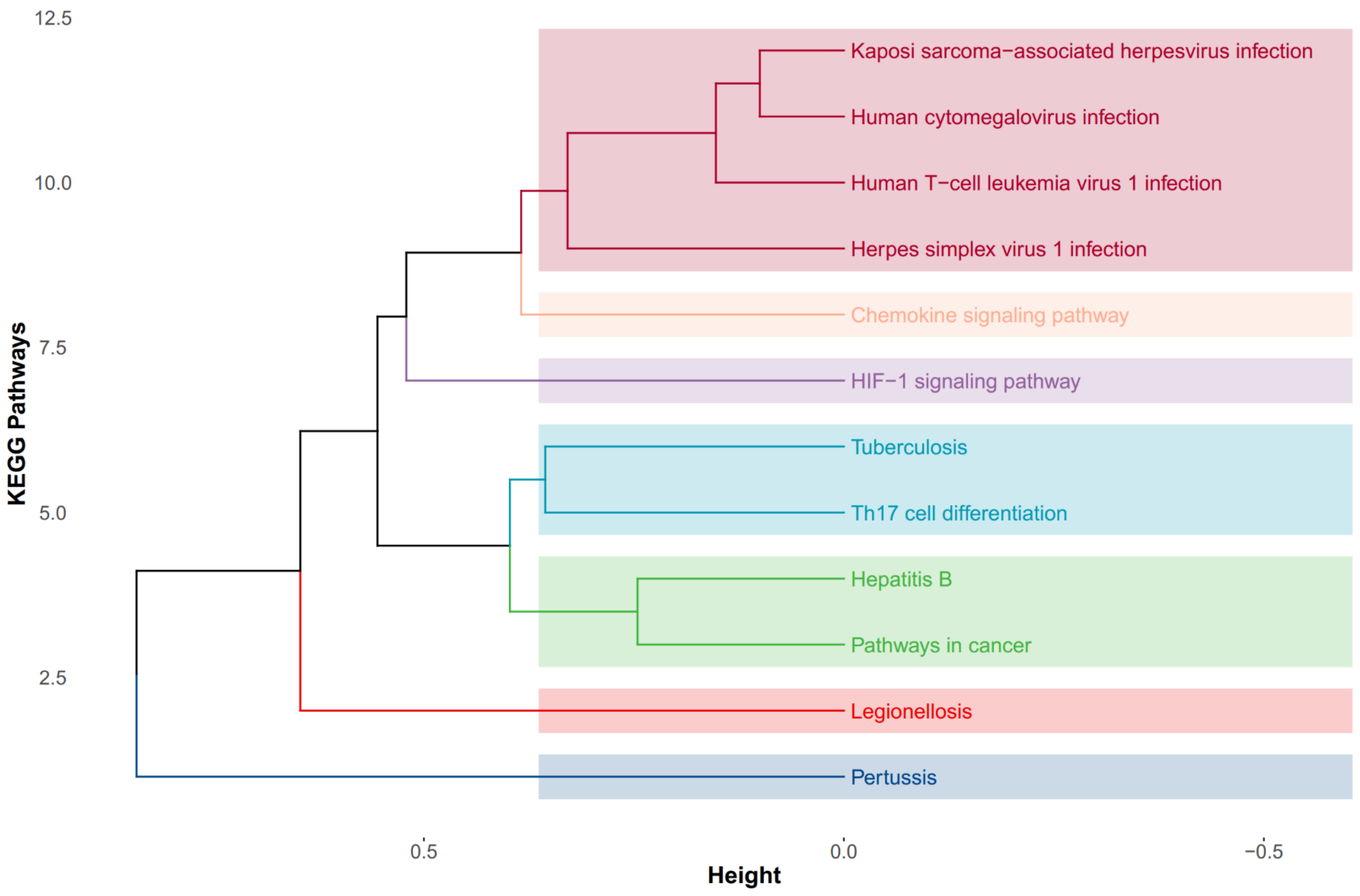

**Figure 2:** Heatmap plot of the interactions between the 67 viral proteins and the 12 final KEGG pathways

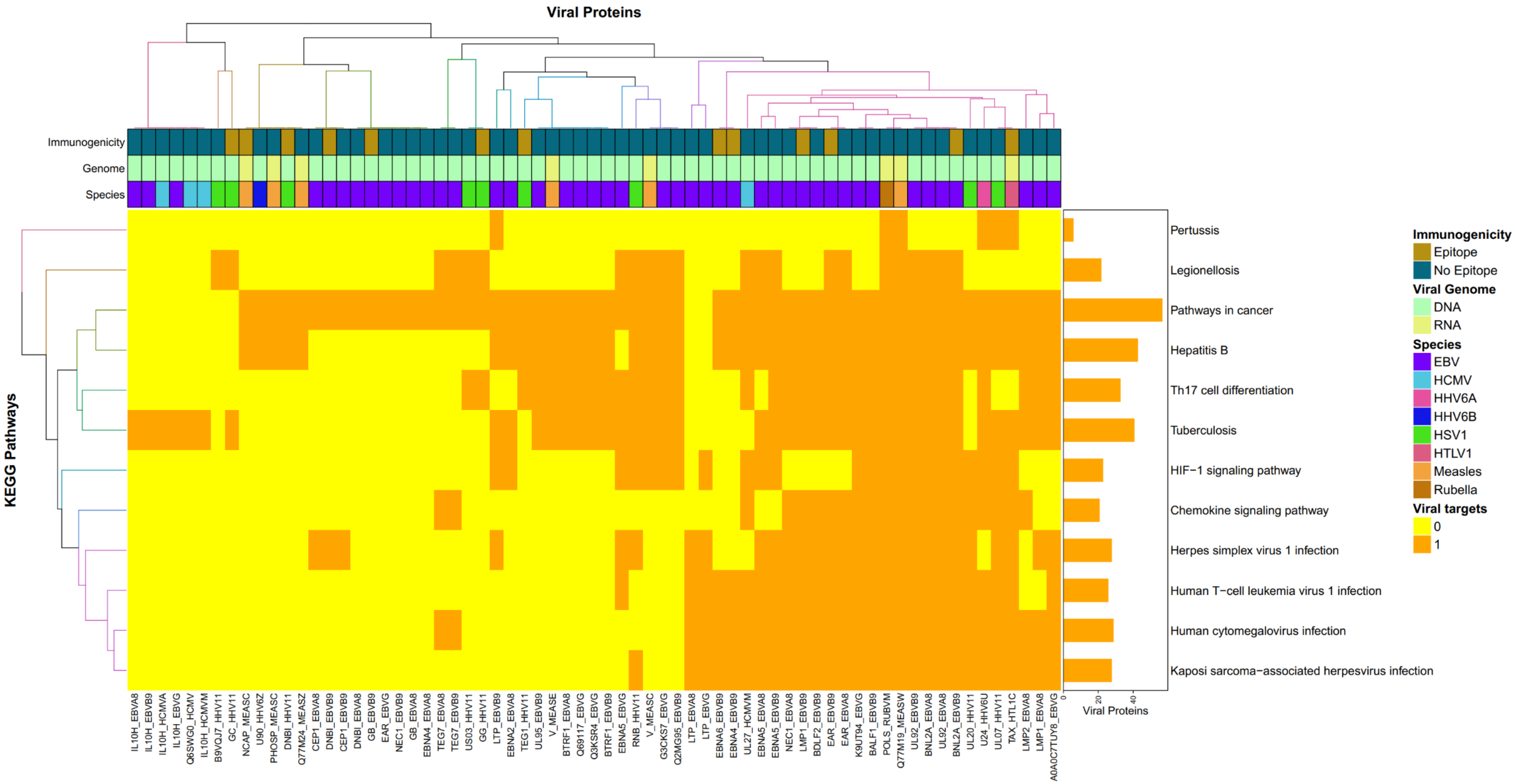
